# The values of ecosystem services inside and outside of protected areas in Eastern and Southern Africa

**DOI:** 10.1101/2023.09.27.559741

**Authors:** Falko T. Buschke, Claudia Capitani

**Affiliations:** European Commission, Joint Research Centre, Ispra, Italy

**Keywords:** Africa, ecosystem services, generalised linear mixed-effect model, nature’s contributions to people, protected areas

## Abstract

Conservation policies often take for granted the importance of protected areas for supplying ecosystem services. The 1^st^ edition of the *State of Protected and Conserved Areas in Eastern and Southern Africa* report contained limited information on ecosystem services, so for the 2^nd^ edition we compared statistically 561 standardised economic values of various types of ecosystem services inside and outside of protected areas. We found that data from local and sub-national case-studies in the Ecosystem Service Valuation Database were biased geographically, highlighting major evidence gaps for most of the region. For well-studied countries (Botswana, Ethiopia, Kenya, South Africa, Tanzania, Uganda), the value of ecosystem services varied considerably across different types of services but were – on average – three to six times higher outside protected areas. This trend was not universal, however, given that opportunities for recreation and tourism tended to be higher within protected areas. Combined, these findings suggest that conservation authorities across Eastern and Southern Africa (1) prioritise ecosystem service valuation studies; (2) expand the focus of ecosystem service policies to include wider landscapes beyond protected area boundaries; and (3) avoid making generic assumptions about ecosystem services by identifying which types of services are most compatible with the broader goals of protection.

## 1. Introduction

Protected areas are valuable for maintaining biodiversity (Watson et al., 2014; Wauchope et al., 2022), and it is often taken for granted that they also enhance ecosystem services, the benefits people obtain from nature. Consequently, global and regional conservation policies regularly link ecosystem services with protected areas. At the global level, Target 3 of the Kunming-Montréal Global Biodiversity Framework aims to conserve areas of particular importance to ecosystem services through protected areas and other effective area-based conservation measures (Convention on Biological Diversity, 2022). In Africa, the *Kigali Call to Action for People and Nature*, which emerged from the inaugural IUCN Africa Protected Areas Congress in 2022, appealed for unprotected areas of importance to ecosystem services to be built into conservation plans (IUCN, 2022).

Despite the implicit acceptance that protected and conserved areas (hereafter protected areas) are important for the continued supply of ecosystem services, empirical evidence from Africa is far from unequivocal. For instance, although the IPBES regional assessment report for Africa (IPBES, 2018) described protected areas as “a driver of positive change” for ecosystem services (page 265), it did not present empirical evidence comparing the ecosystem services inside and outside of protected areas. Similarly, neither of the *State of Protected and Conserved Areas* reports for Central Africa (Doumenge et al., 2021) or Eastern and Southern Africa (IUCN Eastern and Southern Africa Regional Office, 2020) quantified the relative contributions of protected areas to ecosystem services.

Regional protected area policies in Africa would benefit from a transparent scientific foundation when it comes to ecosystem services. Existing empirical studies in Eastern and Southern Africa tend to focus on single protected areas and their immediate surroundings (e.g., Guerbois and Fritz, 2017; Ndayizeye et al., 2020), or on a protected area network within a single country (e.g., Ament et al., 2017; Roux et al., 2020). Other studies quantify the coverage of ecosystem services by a regional protected area network (e.g., Neugarten et al., 2020; Wei et al., 2020), assuming implicitly that ecosystem services are enhanced by protection. Despite this, comparative studies of different types of ecosystems services inside and outside of protected areas remain scarce.

In this study, we evaluate ecosystem services inside and outside of protected areas in Eastern and Southern Africa. Our analysis fed into the 2^nd^ edition of the *State of protected and conserved areas in Eastern and Southern Africa* to guide how ecosystem services within protected areas and surrounding areas ought to be acknowledged, managed, and communicated (IUCN ESARO, 2024). To cover such a large geographic area, we rely on existing ecosystem valuation studies from the Ecosystem Service Valuation Database (ESVD) (Brander et al., 2024, 2023), one of the most comprehensive and standardised collections of ecosystem service valuations studies (Schmidt and Seppelt, 2018). Although the ESVD database is limited to studies that estimated the monetary value of ecosystem services, it has the unique feature that its spatially explicit terrestrial case-studies include information on protection status (Schmidt and Seppelt, 2018). This allowed us to compare statistically the value of different kinds of ecosystem services inside and outside of protected areas in Eastern and Southern Africa.

Our analysis was primarily exploratory, without specific *a priori* hypotheses. However, we started from the position that value varied across different types of ecosystem services, thereby necessitating an analysis that distinguishes between different ecosystem services. In some instances, protection likely shields ecosystems from negative human impacts, enhancing the supply and value of ecosystem services. In other instances, protection may restrict people’s access to ecosystems services, thereby reducing their value. Therefore, we did not have a specific hypothesis about whether the values of ecosystem services were higher or lower in protected areas overall because this would depend on an interaction between protection status and the relative value of different types of ecosystem services. So rather than predicting the specific monetary value of ecosystem services within protected areas, our analysis is intended to provide broad guidance for protected area policies across the region.

## 2. Methods

### 2.1. Study area

This study focused on 26 countries (Figure 1) covered by the 2^nd^ edition of the State of protected and conserved areas in Eastern and Southern Africa (IUCN ESARO, 2024). The 1^st^ edition of the report covered 24 countries in Eastern and Southern Africa (IUCN Eastern and Southern Africa Regional Office, 2020), but the latest edition was expanded to also consider Burundi and the Democratic Republic of Congo (DRC) because both states are members of the East African Community (EAC) and the DRC is member of the Southern African Development Community (SADC).

**Figure 1:**
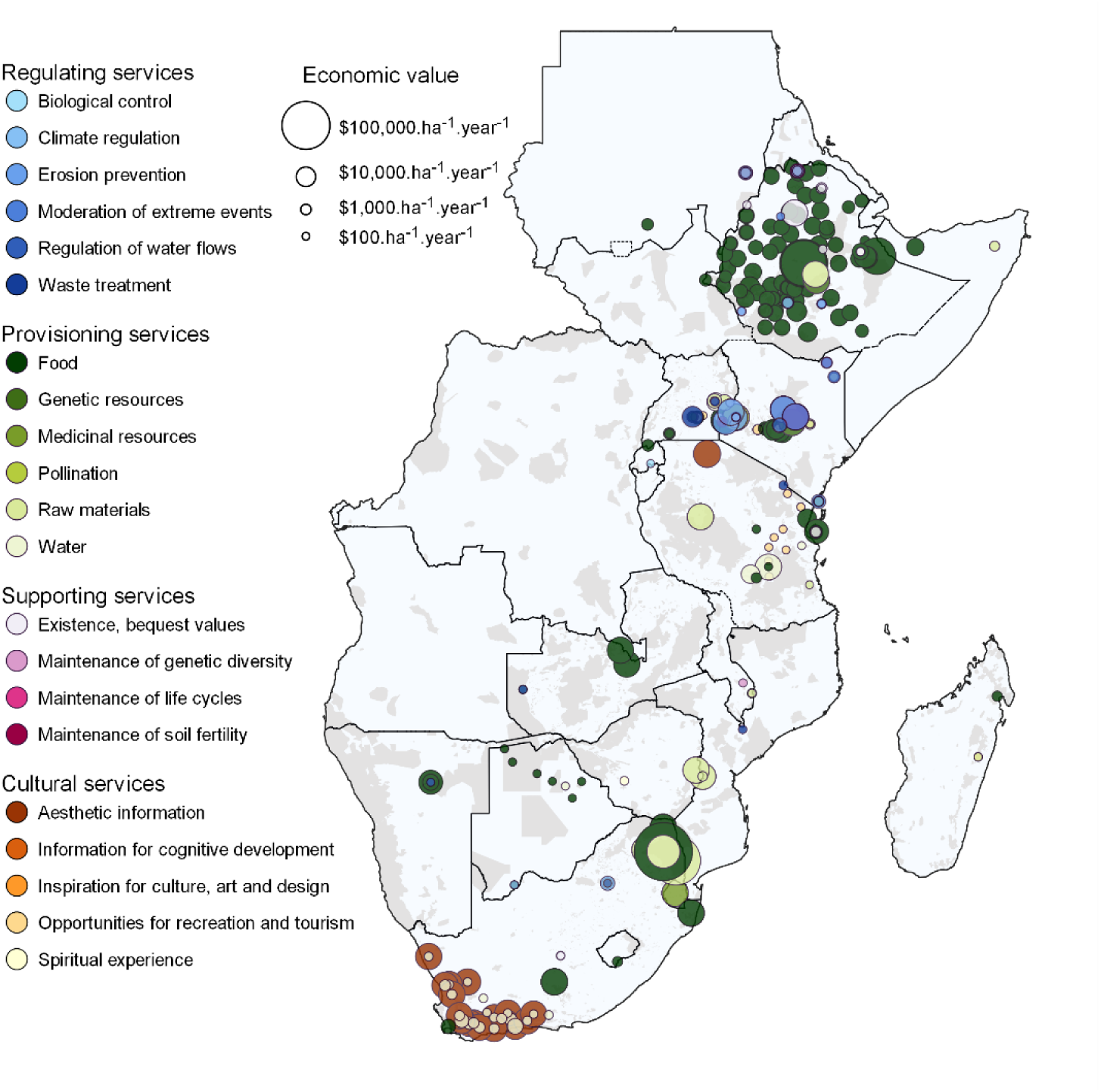
The geographic location of local and sub-national case studies valuing ecosystem services across Eastern and Southern Africa from the Ecosystem Services Valuation Database (Brander et al., 2023). Points are coloured according to the type of ecosystem service and scaled according to the standardised economic value (International $ ha^-1^ year^-1^ for the year 2020). Shaded areas denote protected areas from the World Database for Protected Areas (UNEP-WCMC and IUCN, 2022). The two Indian Ocean Island states of Mauritius and Seychelles are not displayed on this map because there were no local or sub-national ecosystem service valuation case studies for either country.

### 2.2. Data sources and data inclusion rules

Data were from the version of the Ecosystem Service Valuation Database (ESVD) available on 31 May 2023 (Brander et al., 2024, 2023), which we filtered to a relatively balanced subset for analyses based on the attributes reported for each record (Figure 2). This global database is comprised of approximately 10,000 estimated values of ecosystem services from more than 1,100 case studies. Values in the database are derived from published literature sources (both peer-reviewed and grey literature), which meet criteria of minimum information on the year of assessment, geographic locality, ecosystem or biome identity, type of ecosystem service, valuation metric, and valuation method (Brander et al., 2024).

**Figure 2:**
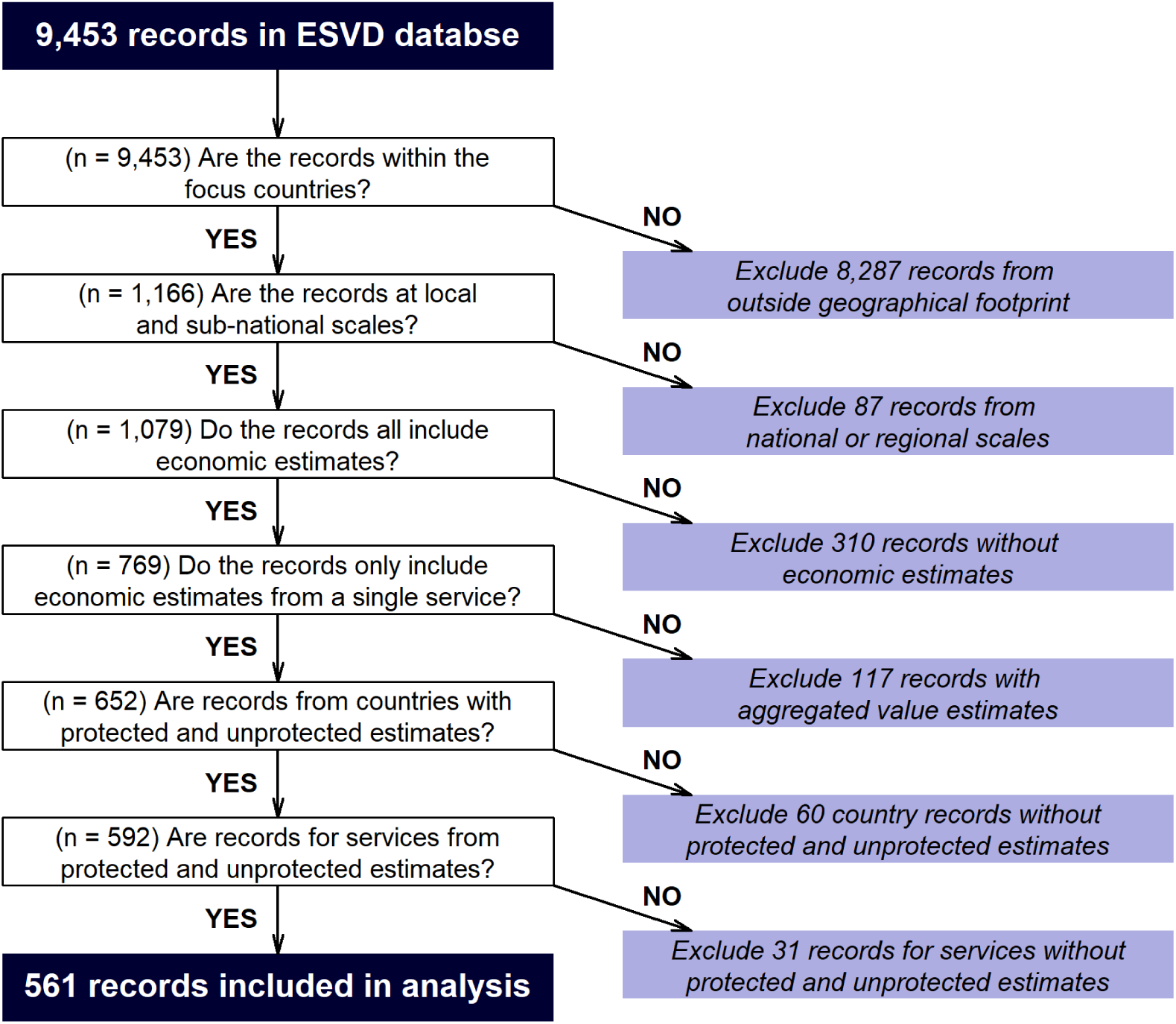
A schematic decision-tree of the rules used to include data from the Ecosystem Service Valuation Database into a generalised mixed effects linear model.

We filtered the global dataset to the 26 focal countries of this study, resulting in 1,166 independent value estimates. We only included estimates from local (n = 866) and sub-national (n = 213) assessments. Each estimate included a standardised value of ecosystems services in International $ha^-1^year^-1^ for the year 2020, which is the value of the inflation-adjusted US$ in the United States in terms of purchasing power (Brander et al., 2024). After excluding cases where standardised economic value was either unreported or reported as zero, 769 estimates remained (these are shown in Figure 1).

Each record in ESVD includes metadata on protection status. This information includes three levels of protection: “*No protection*” (n = 187); “*Partially Protected*” (n = 352); and “*Protected*” (n = 106), which we aggregated into two levels by combining “*Partially Protected*”, and “*Protected*”. We also assumed that records without protection information (n = 124) were unprotected, so we combine these into a single “*Unprotected”* category.

The ESVD identifies the specific type of ecosystem service for each estimate of value (Figure 1), but in some instances, estimates represent the aggregated value of multiple ecosystem services. The database guidelines advise against disaggregating values from multiple services, so we excluded all estimates based on multiple ecosystem services (n = 117).

To balance the data included in the statistical analysis, we only included data from countries that had estimates from both protected and unprotected sites (n = 592). This meant that we excluded data from Madagascar (n = 7 protected sites), Malawi (n = 3 protected sites), Mozambique (n = 5 unprotected sites), Namibia, (n = 10 protected sites), Rwanda (n = 1 protected site), Sudan (n = 21 protected sites), Zambia (n = 5 protected sites), and Zimbabwe (n = 8 unprotected sites). Similarly, we excluded types of ecosystem services that did not include estimates from both protected and unprotected sites. This resulted in the exclusion of “*unreported”* ecosystem service types (n = 12 unprotected sites); “*aesthetic information”* (n = 1 unprotected site); “*inspiration for culture, art, and design”* (n = 1 protected site); “*maintenance of genetic diversity”* (n = 2 protected sites); “*maintenance of life cycles”* (n = 1 protected site); and “*regulation of water flows”* (n = 2 protected sites). After these exclusions, Somalia no longer included estimates from unprotected sites, so we excluded the remaining estimates from protected sites from this country (n = 12). The final dataset used for statistical modelling included 561 estimates of 12 ecosystem service types from both protected and unprotected sites in six countries (Botswana, Ethiopia, Kenya, South Africa, Tanzania, and Uganda). These estimates were relatively balanced across protection levels and types of ecosystem services (Figure S2).

### 2.3. Model structure

Our aim was to examine whether protected areas are associated with higher values of ecosystem services, while recognising that protection may be more effective at conserving certain types of ecosystem services than others. As such, we included protection level and ecosystem type, and their interactions, as categorical fixed effects in our model. Moreover, we anticipated that the relationship between protected areas and ecosystem services was possibly affected by several unmeasured spatially autocorrelated variables – such as climate, biogeography, governance, or management – which would have been too complex to model explicitly. Therefore, we included countries as random effects in the mixed effects model.

The distribution of values of ecosystems services (International $ ha^-1^ year^-1^ for the year 2020) were strongly right-skewed and dominated by many low values and few high values. Therefore, we initially considered two modelling approaches: (1) a log-linked Gamma generalised linear mixed effect model (*Gamma-glme*) using the `glmer` function in the `lme4` package (Bates et al., 2015) in R version 4.1.3 (R Core team, 2023), and (2) a linear mixed effects model with a log-transformed response variable (*log-lme:* `lmer` function in `lme4` package). Model diagnostics suggested that neither modelling approach was clearly superior (Appendix 1, Figure S1). So we favoured the *Gamma-glme* because the model residuals were distributed more normally than the *log-lme* (Figure S1 a,b) and the model predictions are more intuitive (i.e., *Gamma- glme* models the log of the expected mean value of ecosystem services, whereas the *log-lme* models the expectation of the log-transformed mean value of ecosystem services: Appendix 1). Our final model structure was:

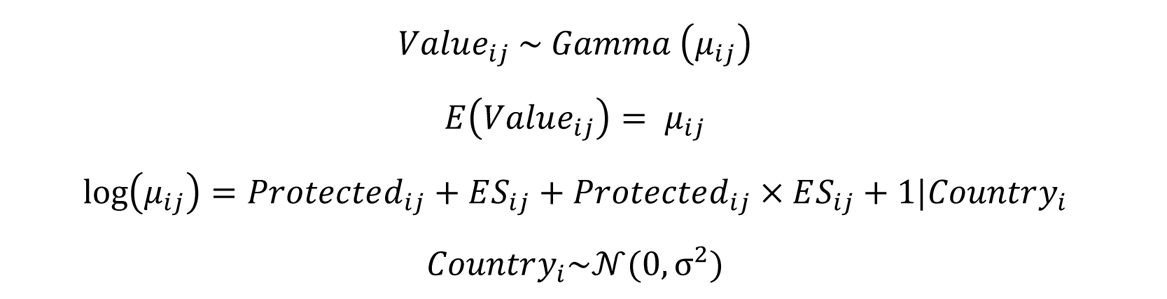

Where *Value*_ijk_ is the estimated values (International $ ha^-1^ year^-1^ for the year 2020) of ecosystem observation *j* in country , which we assume are random variables from a Gamma distribution with an expected (*E*) mean value of *μ*_ijk_. *Protected*_ijk_ (two levels of protection, with “*Unprotected”* as the baseline) and *ES*_ijk_ (12 levels of ecosystem service type, with “*Food”* as the baseline) were included as categorical predictor variables, along with their two-way interaction. *Country*_i_ is the random intercept in the model, which is assumed to be normally distributed with a mean of zero and a variance of *σ*^2^.

Alongside this full model, we also modelled three subset models by (i) only including protection as a predictor, (ii) only including ecosystem service type as a predictor, (iii) modelling protection level and ecosystem service type additively without an interaction term. These subset models were compared using a likelihood-ratio *X*^2^ test (`anova` command in `stats` package). Given the relatively large dataset – 561 data points – we used a conservative threshold (*α* = 0.01) to assess whether regression coefficients differed significantly from zero.

As a supplementary analysis (Appendix 2), we also analysed a second model formulation, which also included biome as a random effect in the model. Data were unevenly balanced across countries, biomes, or types of ecosystem service (Figures S2 and S3). This led to convergence issues for the more complex model formulation, and likely overfitted the model to the data. Despite these issues, the results from the more complex formulation were consistent with those from the simpler model (Table S1): the main statistical effects were the same, and the majority of the estimated coefficients had the same effect (i.e. sign) and statistical significance across both model formulations.

## 3. Results

Estimates of ecosystem service values in the ESVD were clearly biased geographically (Figure 1). Eight of the 26 countries (30%) in Eastern and Southern Africa had no case study data, and a further seven countries had fewer than 10 value estimates. Therefore, our statistical model should be interpreted cautiously; especially in parts of the region that are poorly represented in the ESVD.

The log-linked Gamma generalised linear mixed effects model that included both protection level, ecosystem service type and their interaction provided the best fit to the empirical data from the ESVD (Table 1). This model formulation had the lowest AIC (7734.9) and highest log-likelihood (-3841.5) compared to simpler model formulations. This confirms that the variation in the standardised economic value of ecosystem services is related statistically to protection status, the type of ecosystem service and their interaction.

**Table 1:** An incremental comparison of candidate model formulations of increasing complexity using a likelihood-ratio *X*^2^ test. Models are ordered based on decreasing Akaike’s Information Criterion (AIC), which corresponds with the addition of fixed effects – and their interaction – to the model.

| Fixed effects | AIC | log-likelihood | $\chi^2$ (degrees of freedom)* | P-value* |
| --- | --- | --- | --- | --- |
| Protection | 7822.2 | -3907.1 | - |  |
| Ecosystem service type | 7790.5 | -3881.2 | 51.749 (10) | < 0.001 |
| Protection + Ecosystem service type | 7757.2 | -3863.6 | 35.296 (1) | < 0.001 |
| Protection + Ecosystem service type + (Protection x Ecosystem service type) | 7734.9 | -3841.5 | 44.255 (11) | < 0.001 |

\* Values are based on a pair-wise comparisons with the preceding model

The variance explained by country identity as a random effect to the full model accounted for 85.84 % of the total random variance (Variance_country_ = 29.78 ± 5.5 sd; Variance_residualS_ = 4.91 ± 2.2 sd). The average value of ecosystem services increased sequentially across Botswana, Tanzania, Kenya, Ethiopia, Uganda, and South Africa (Figure 3). It remains uncertain whether this is due to differences between the countries *per se* (e.g., governance or management differences) or due to underlying spatially autocorrelated environmental variables (e.g. climate, ecosystem type, or biogeography). Nevertheless, it cautions against assuming that the standardised economic value from one part of the region is transferable to another.

**Figure 3:**
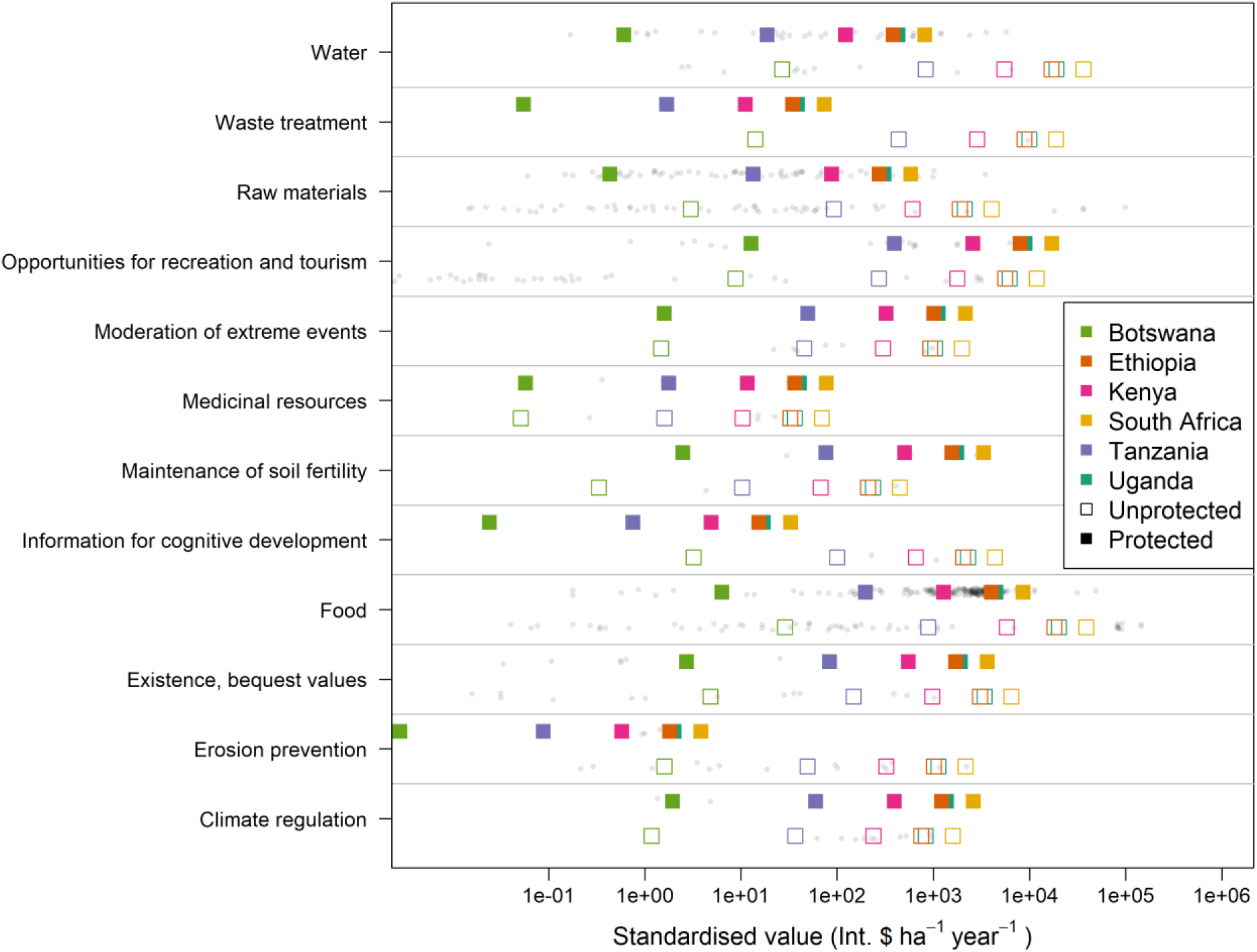
The standardised value (International $.ha^-1^.year^-1^) of different types of ecosystem services in protected and unprotected areas. Grey points show empirical values from 561 case studies. Coloured squares show predictions from a log-linked Gamma generalised linear mixed effects models with six countries included as random factors. The horizontal axis is log-scaled.

Model coefficients suggest that the standardised value of ecosystem services outside of protected areas was three to six times higher than the value inside protected areas (*e*^−1.505(±0.31)^ = 0.22 (0.16 − 0.30): Table 2). This large effect size was also statistically significant (*t =* -4.89, P < 0.001). Standardised economic value also varied across the type of ecosystem service (Table 2). “*Food”* had the highest value on average and was set as the baseline category in the regression model (Table 2, Figure 3). The values of “*existence, bequest values”, “information for cognitive development”, “opportunities for recreation and tourism”, “waste treatment”,* and *“water”* were statistically similar to the value of “*food”* (each of these categories had negative coefficients suggesting that they were less valuable on average, but these coefficients did not differ significantly from zero at the conservative threshold of 0.01: Table 2). By contrast, the values of *“climate regulation”* (2.1 – 8.1 % the value of food), *“erosion prevention”* (3.0 – 10.3 % the value of food) *“maintenance of soil fertility”* (0.3 – 4.5 % the value of food), *“medicinal resources”* (2.3 – 11.3 % the value of food), *“moderation of extreme events”* (2.3 – 11.3 % the value of food ), and *“raw materials”* (6.1 – 17.7 % the value of food) were all considerably less than *“food”* (Table 2; Figure 3).

**Table 2:** Model coefficients of a log-linked Gamma generalised linear mixed effects model of the standardised value of ecosystem services relative to protected status, the type of ecosystem service and their interaction. Given the log-link in the model, exponents of the estimated coefficients reflect the statistical effect on the value of ecosystem services. Statistically significant coefficients at the conservative threshold of *α* = 0.01 are shown as P-values in bold font.

| Fixed effect | Additive terms |  |  | Interaction terms |  |  |
| --- | --- | --- | --- | --- | --- | --- |
|  | Coefficient (SE) | t-value | P-value | Coefficient (SE) | t-value | P-value |
| Protection (Unprotected = 0) |  |  |  |  |  |  |
| *Protected* | -1.505 (0.31) | -4.89 | **< 0.001** |  |  |  |
| Ecosystem service type (Food = 0) |  |  |  | **Interaction with protection** |  |  |
| *Climate regulation* | -3.189 (0.67) | -4.75 | **< 0.001** | 1.999 (1.27) | 1.57 | 0.116 |
| *Erosion prevention* | -2.887 (0.61) | -4.71 | **< 0.001** | -4.824 (1.25) | -3.87 | **< 0.001** |
| *Existence, bequest values* | -1.780 (0.71) | -2.51 | 0.012 | 0.928 (1.01) | 0.92 | 0.360 |
| *Information for cognitive development* | -2.183 (1.38) | -1.59 | 0.113 | -3.386 (2.33) | -1.46 | 0.146 |
| *Maintenance of soil fertility* | -4.459 (1.36) | -3.28 | **0.001** | 3.519 (1.75) | 2.02 | 0.044 |
| *Medicinal resources* | -6.324 (1.0) | -6.32 | **< 0.001** | 1.615 (1.47) | 1.10 | 0.273 |
| *Moderation of extreme events* | -2.969 (0.79) | -3.74 | **< 0.001** | 1.590 (2.04) | 0.78 | 0.435 |
| *Opportunities for recreation and tourism* | -1.181 (0.55) | -2.15 | 0.032 | 1.874 (0.61) | 3.08 | **0.002** |
| *Raw materials* | -2.260 (0.53) | -4.28 | **< 0.001** | -0.425 (0.56) | -0.76 | 0.446 |
| *Waste treatment* | -0.714 (1.96) | -0.37 | 0.715 | -4.045 (2.68) | -1.51 | 0.131 |
| *Water* | -0.062 (0.76) | -0.08 | 0.935 | -2.288 (0.93) | -2.46 | 0.014 |

The model’s interaction coefficients suggested that protection status affected the value of ecosystem services differently (Table 2). Specifically, the value of *“erosion prevention*” in protected areas was more than an order of magnitude less than the value of the same service outside of protected areas. In comparison, the average value of *“opportunities for recreation and tourism”* inside protected areas was roughly 45% higher than in unprotected areas (Table 2; Figure 3).

## 4. Discussion

This study set out to analyse the value of different types of ecosystem services inside and outside of protected areas in Eastern and South Africa. Our analysis was designed to inform the 2^nd^ edition of the *State of protected and conserved areas in Eastern and Southern Africa* (IUCN ESARO, 2024). However, the message most relevant to policy probably stems from what our analysis failed to show. The uneven geographic spread of records in the ESVD database highlights how little is known about the value of ecosystem services across most of the region and suggests that collecting representative data should be an urgent policy priority across the region.

A recent study by Duncanson et al., (2023) quantified the added value of protected areas for preserving carbon stocks globally. Five Eastern and Southern African countries (DRC, Mozambique, Tanzania, Zambia, and Madagascar) were amongst the top 20 countries globally in terms of the additionality of carbon storage by protected areas. Yet despite these countries being important for global climate mitigation, only Tanzania had sufficient ecosystem valuation case studies to inform our comparative analysis. Data gaps like these risk overlooking much of nature’s value, leading to its inadvertent devaluation in decision-making contexts (Costanza et al., 2014). Such disregard is unfortunately already a common policy outcome even when valuation data do exist (IPBES, 2022; Laurans et al., 2013). Although we did not expressly set out to compare the value of ecosystem services across countries in Eastern and Southern Africa, the random effect in our statistical model showed how country identity accounted for 85% of the variation in ecosystem service value that could not be attributed to the protection status or type of ecosystem service. This suggests that filling data gaps in one country by extrapolating the value of ecosystem services to another is not advisable.

The data we could access suggested that the value of ecosystem services was on average five times higher outside of protected area than within them. This relationship is unlikely causal and does not imply that protected area expansion will reduce the value of ecosystem services. The monetary value of ecosystem services is not just a function of the aggregate supply of these services by nature, but also the aggregate demand by people (Dasgupta, 2021). Aggregate demand is typically determined by human population size, their *per capita* consumption, and the efficiency with which nature’s contributions are consumed (Dasgupta, 2021). All three of these variables tend to be higher outside of protected areas. In our analysis, this is most clearly illustrated by “*food”* ecosystem services, which had the highest value of the service types considered in our model. Even though cropland is expanding into protected areas globally, in Southern and Eastern Africa expansion rates are slower than the rates in unprotected areas (Meng et al., 2023). It stands to reason that the value of food services will be higher outside of protected areas. That said, expanding protected areas would not necessarily reduce cultivated area and the value of food services. Instead, protected area expansion may redirect cultivation elsewhere in the landscape (Ford et al., 2020; Waldron et al., 2020). So, our results should not be misconstrued that protection reduces the value of ecosystem services.

Our results imply that policies specifically targeting ecosystem services should expand their focus beyond the boundaries of protected areas, often in areas already occupied by human settlements. This is consistent with global calls for integrated management of “working” or “shared” landscapes (Kremen and Merenlender, 2018; Obura et al., 2021); regional programmes, like the European Commission’s *NaturAfrica* initiative (European Commission, 2021); and the recognition of other effective area-based conservation measures (Dudley et al., 2018; Gurney et al., 2021). Conservation interventions in unprotected human-dominated landscapes rely on many of the same management approaches as in protected areas: such as clearly defined resource boundaries, the need for monitoring, restricted resource use, and sanctions for rule-breaking (Mahajan et al., 2021). However, across Eastern and Southern Africa these kinds of landscape-wide efforts often struggle for legitimacy and face challenges around governance, power dynamics and environmental justice (Bourgeois et al., 2023; Cassidy, 2021; Kicheleri et al., 2021). There exists a unique tension between the concentration of decision-making power in formal institutions (e.g., conservation authorities) and the local communities who resist these trends (Kicheleri et al., 2021; Nelson et al., 2021). Therefore, conservation authorities should not presuppose that existing protected area policies will transfer smoothly in unprotected landscapes and should plan accordingly.

The significance of the unprotected landscapes for the supply of ecosystem services should not deprioritise these services within the boundaries of protected areas. On the contrary, our results support more nuanced inclusion of ecosystem services in protected area policies and strategies. This is demonstrated by the interaction term in our regression model, which showed, for example, how “*opportunities for recreation and tourism*” were 45% higher in protected areas on average. Globally, the consumer surplus generated by protected area visitation dwarfs spending on protected areas (Balmford et al., 2015). The African Union’s post-pandemic *Green Recovery Action Plan* recognises this opportunity, calling for the immediate support of ecotourism within the context of improving the management of protected areas (African Union, 2021). Protected area authorities could consider management interventions that enhance tourism opportunities because accessibility is a major determinant of tourist visitation to, as well as within, protected areas (Arbieu et al., 2018; Balmford et al., 2015). Such interventions should be prudent though, because unchecked tourism may exacerbate ecological degradation (Dasgupta, 2021; Hunter and Shaw, 2005).

In this study, we were careful not to overinterpret the results by making context-specific policy recommendations. Similarly, we caution others from using these results to draw conclusions about specific countries, protected areas, or types of ecosystems services. Instead, we urge researchers and protected area authorities to supplement these results with local information to overcome three main caveats from this study. First, our results are based on data from only one database. Although the ESVD database has many features that make it suitable for this type of comparative analysis (Brander et al., 2024; Schmidt and Seppelt, 2018), it likely underestimates the full tangible and intangible value of ecosystems services. Second, our study does not consider the full range of ecological dynamics that give rise to ecosystem services. Specifically, our analyses did not distinguish trade-offs between ecosystems services (Howe et al., 2014), or situations where the downstream benefits from unprotected ecosystem services may have originated from within protected sites (e.g., freshwater provisioning, sediment retention, pollinators). Third, our study does not consider the costs of ecosystem services (i.e., disservices), nor the potential mismatches between the beneficiaries of ecosystem services and those who bear the costs from living in proximity with nature (Swemmer et al., 2017). Therefore, we are hopeful that our study serves as the skeleton for a larger body of knowledge about the true value of ecosystem services in the region.

In conclusion, protected areas remain a cornerstone of conservation policy. But our study, which is not without its caveats, advises against the broadbrush association between protected areas and the value of ecosystem services. In general, evidence for such claims is sparse, emphasising the urgent need for ecosystem service valuation studies that (i) address existing geographic biases in data; (ii) evaluate values beyond the narrow scope of monetary valuations studies (e.g., IPBES, 2022); (iii) include assessments of the marine realm, which was completely overlooked by our analyses. The limited data that do exist, however, suggest that ecosystem services provide considerable value beyond the boundaries of protected areas, probably because this is where nature’s services are more accessible. Therefore, policies targeting ecosystem services need to look beyond protected areas to preserve nature’s contributions in unprotected working landscapes. Within protected areas, authorities should avoid generic assumptions about ecosystem services. Tailored interventions require a more nuanced, evidence-based approach. If ecosystem services are a policy priority for protected area authorities, then they should invest resources and attention into identifying the specific types of services most compatible with the broader goals of protection as expressed in individual management plans.

## Acknowledgements

We thank Mark Gerrard and Nuredin Juhar for the collaboration on the 2^nd^ edition of the *State of Protected and Conserved Areas in Eastern and Southern Africa* report.

## Data availability

The data used in this study are available from the Ecosystem Services Valuation Database (https://www.esvd.net/) and the R-scripts to replicate our analysis are available on GitHub (https://github.com/falko-buschke/SOPACA/) and will be permanently deposited on Zenodo after peer-review.

## Equity, Diversity, and Inclusion statement

The authors acknowledge that this study focuses on a region underrepresented in the scientific literature. While this specific manuscript does not include contributions by authors from the region, it only forms one portion of a chapter for the 2^nd^ edition of the *State of Protected and Conserved Areas in Eastern and Southern Africa* report co- authored with researchers from South Africa and Ethiopia (named in the acknowledgments section of this manuscript). Other chapters of the report are authored primarily by researchers from within the region, so this manuscript forms part of a larger representative research process.

## Supporting Information

### Appendix 1: Model diagnostics

In the main manuscript, we present the results from a log-linked Gamma generalised linear mixed effect model (*Gamma-glme*), which we favoured over a linear mixed effects model with a log-transformed response variable (*log-lme*).

Model diagnostics suggested that neither approach was clearly superior. The residuals from the *Gamma-glme* model (Figure S1a) were distributed more normally than those from the *log-lme* model (Figure S1b), thus favouring the former model formulation. However, the residuals from the *Gamma-glme* model were biased downwards (mean residuals were negative: Figure S1c) compared to those from the *log-lme* model (Figure S1d). The modes from the random intercepts were approximately normal for both the *Gamma-glme* (Figure S1e) and the *log-lme* (Figure S1f), though in the former model intercepts for Botswana were lower than expected from normally distributed random effects.

A pertinent difference between the two modelling approaches, however, is how they model the response variable. The *Gamma-glme* model estimated the log of the expected mean value of ecosystem services, *log*(*E*[*Value*]), whereas the *log-lme* models the expected mean of the log-transformed value of ecosystem services, *E*[*log*(*Value*)]. These estimates are not the same. Given that the distribution of ecosystem service values is strongly right-skewed, modelling log-transformed response variables downplays ecosystems services with exceptionally high value. Therefore, we used the *Gamma-glme* model for the rest of our study.

### Appendix 2: Incorporating information on ecological biome in the mixed effects model

In the main manuscript, we presented a mixed effects model that included country identity as a random factor. We also examined a second model formulation, that also included biome identity as a random factor, which we present here.

The subset of the ESVD database used for our model included ecological information on the *biome* (13 categories), *ecosystem* (13 categories), and *ecozone* (23 categories) for each record. We only considered *biome* identity in our model because (a) the *ecosystem* data were not reported for two-thirds of the records (368 of 561 records), and (b) 23 categories of *ecozone* were too many for our model and led to convergence issues.

We included biome as a random factor in the model using the following model structure:

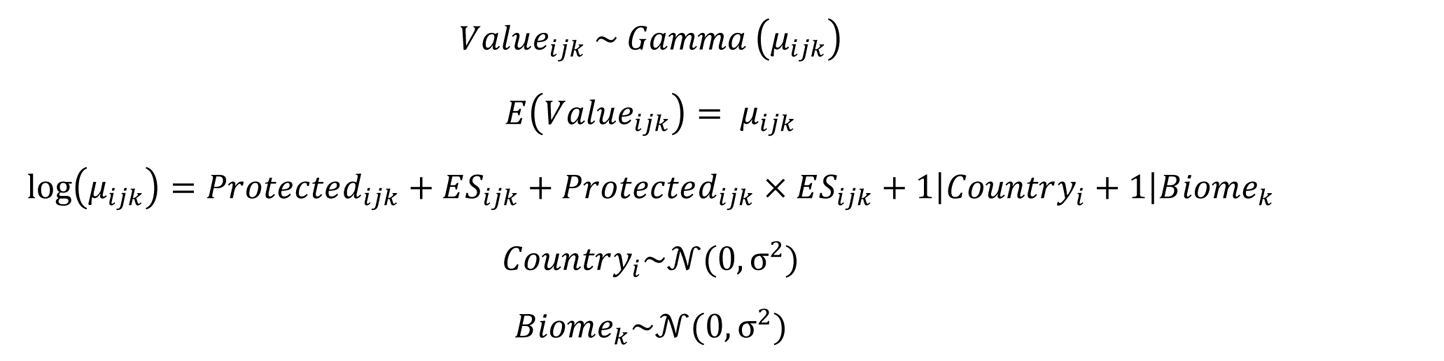

Where *Value*_ijk_ is the estimated value (International $ ha^-1^ year^-1^ for the year 2020) of ecosystem observation *j* in country i and biome *k* with an expected (*E*) mean value of *μ*_ijk_. *Protected*_ijk_ (two levels of protection, with “*Unprotected”* as the baseline) and *ES*_ijk_ (12 levels of ecosystem service type, with “*Food”* as the baseline) were included as categorical predictor variables, along with their two-way interaction. *Country*_i_ and *Biome*_*k*_ are random intercepts in the model, which are assumed to be normally distributed with a mean of zero and a variance of *σ*^2^.

Our data inclusion rules were based on balancing records across ecosystem service types, protection levels, and countries (Figure S2). The consequence of this decision was that records were weakly balanced between countries, ecosystem services, and biomes (Figure S3), which led to convergence issues in the generalised mixed effects models. Specifically, our dataset included information on 56.9% of all combinations of country and type of ecosystem service (41 of 72 combinations), but only 42.3% of combinations of country and biome (33 of 78 combinations) and 37.2% of combinations of biome and type of ecosystem service (58 of 156 combinations). Although these convergence issues could be resolved by using a different model optimiser (i.e. bound optimization by quadratic approximation, BOBYQA) and using longer runs to lower convergence tolerances, the risk of overfitting the model remained. Therefore, while we present the results of the model with biome as a random factor in this appendix, we preserve the more conservative outputs from the simpler model for the main manuscript.

The variance explained by country identity as a random effect to the full model accounted for 49.07 % of the total random variance, whereas the variance explained by biome accounted for 46.49% of the random variance (Variance_country_ = 38.98 ± 6.2 sd; Variance_biome_ = 36.93 ± 6.1 sd; Variance_residuals_ = 3.53 ± 1.9 sd). Together country and biome identity accounted for 95.56% of the random variance in the model, which may signal overfitting.

In general, the model coefficients from the full model with two random effects were consistent with those of the simpler model presented in the main manuscript (Table S1). When considering biome identity, the average estimated value of ecosystem services outside of protected areas was 10-times higher compared to protected areas (*e*^−2.420^ = 0.09: Table S1), suggesting that the results reported in the main manuscript may possible be conservative.

Six of the eleven coefficients for the type of ecosystem service indicated comparable outcomes as the simpler model in the main manuscript; that is the sign and statistical significance of the coefficients were the same across the two models. For three coefficients (“*Existence, bequest values*”, “*moderation of extreme events*”, and “*opportunities for recreation and tourism*”), the signs of the coefficients were the same across the two models, though their statistical significance differed. By contrast, coefficients for “*climate regulation*”, which were significantly positive in the simple model presented in the main manuscript, were marginally positive (without being statistically significant) in the more complex model that accounted for differences in biomes. As with the model in the main manuscript, “*Food”* generally had the highest average value of the different types of ecosystems services. However, in the model that considered biome identity, the value of water services was nearly eight-times higher than that of food (775%). This is likely because the value of water services is relatively higher in arid biomes, where the overall values of ecosystem services tends to be lower on average.

Six of the eleven interaction coefficients remained the same (i.e., same sign and significance) when considering biome in the model, and a further two coefficients switched signs without being statistically distinguishable from zero in either model formulation. In the case of interaction coefficient for “*erosion prevention*”, which was significantly negative in the simpler model, considering the biome in the model meant that the coefficient was no longer statistically significant. By contrast, interaction coefficients for “*Medicinal resources*” were significantly positive when considering biome difference, compared to being positive but not statistically different in the simpler model. The interaction coefficient for “*Raw materials*” changed sign in a statistically significant way when considering biome identity, suggesting that the value of raw materials inside protected areas are 71.3% higher than unprotected areas.

In summary, incorporating biome as a random factor in the model would not overturn the results reported in the main manuscript in a way that would affect the interpretation or conclusions of the study. However, they do emphasise the need for a nuanced approach to evaluating the value of ecosystem service, especially when drawing conclusions from imperfect observational data.

**Figure S1:**
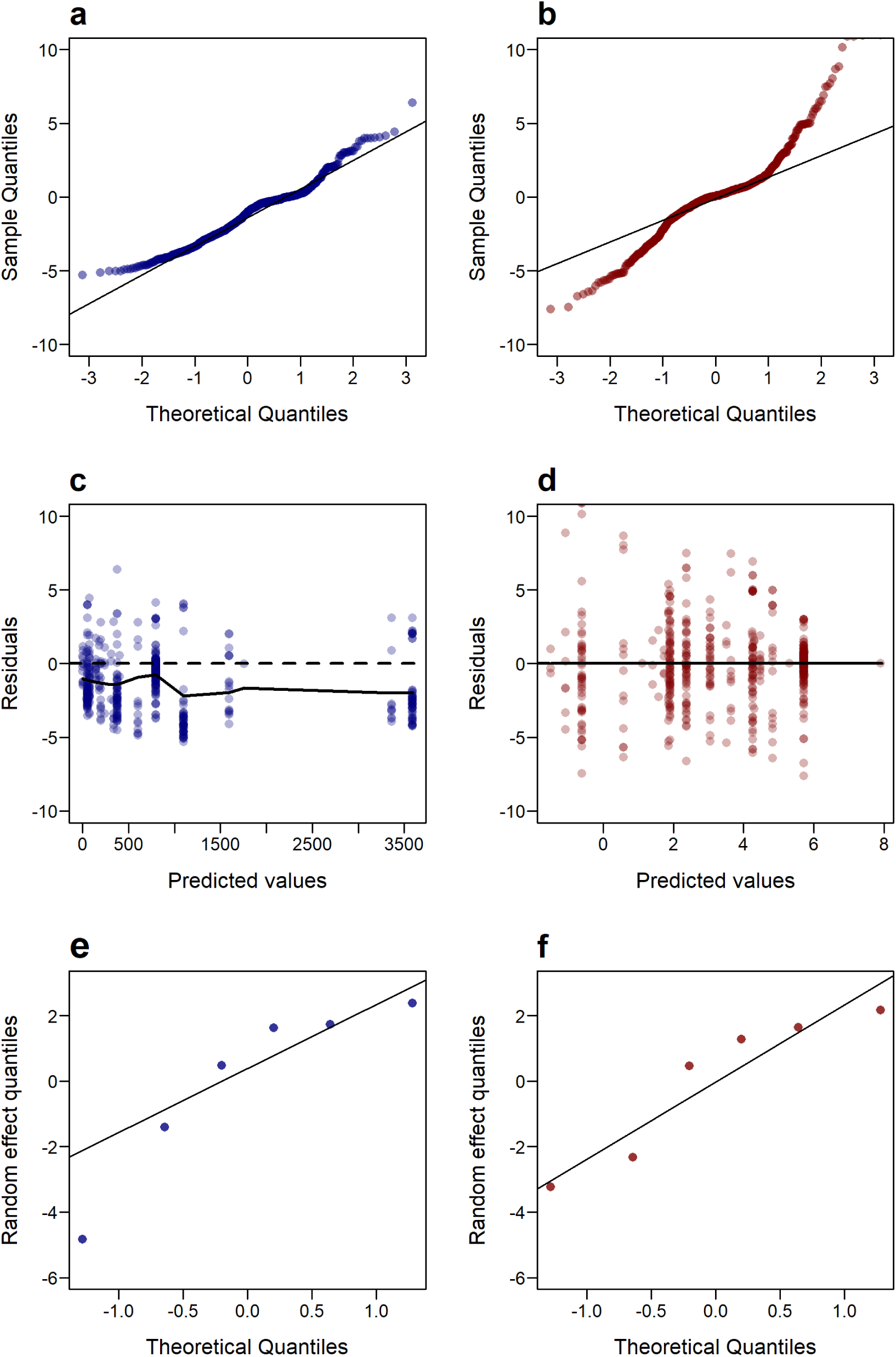
Diagnostic plots comparing a log-linked generalised linear mixed effect model to linear mixed effects model with a log-transformed response variable. Q-Q normal plots for the model residuals (**a** and **b**), the residual plot relative to model predictions (**c** and **d**), Q-Q normal plots for the intercepts of the country identity as a random effect in the model (**e** and **f**).

**Figure S2:**
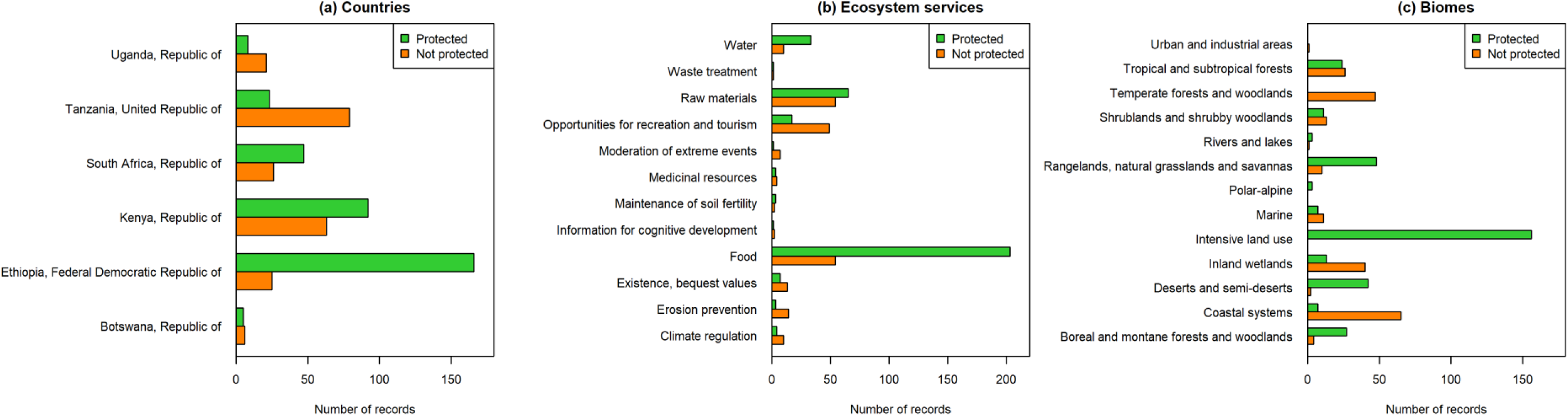
Comparing the number of records used in our analyses for ecosystem service values inside and outside of protected areas for (a) countries, (b) the types of ecosystem service, and (c) the biome of the study.

**Figure S3:**
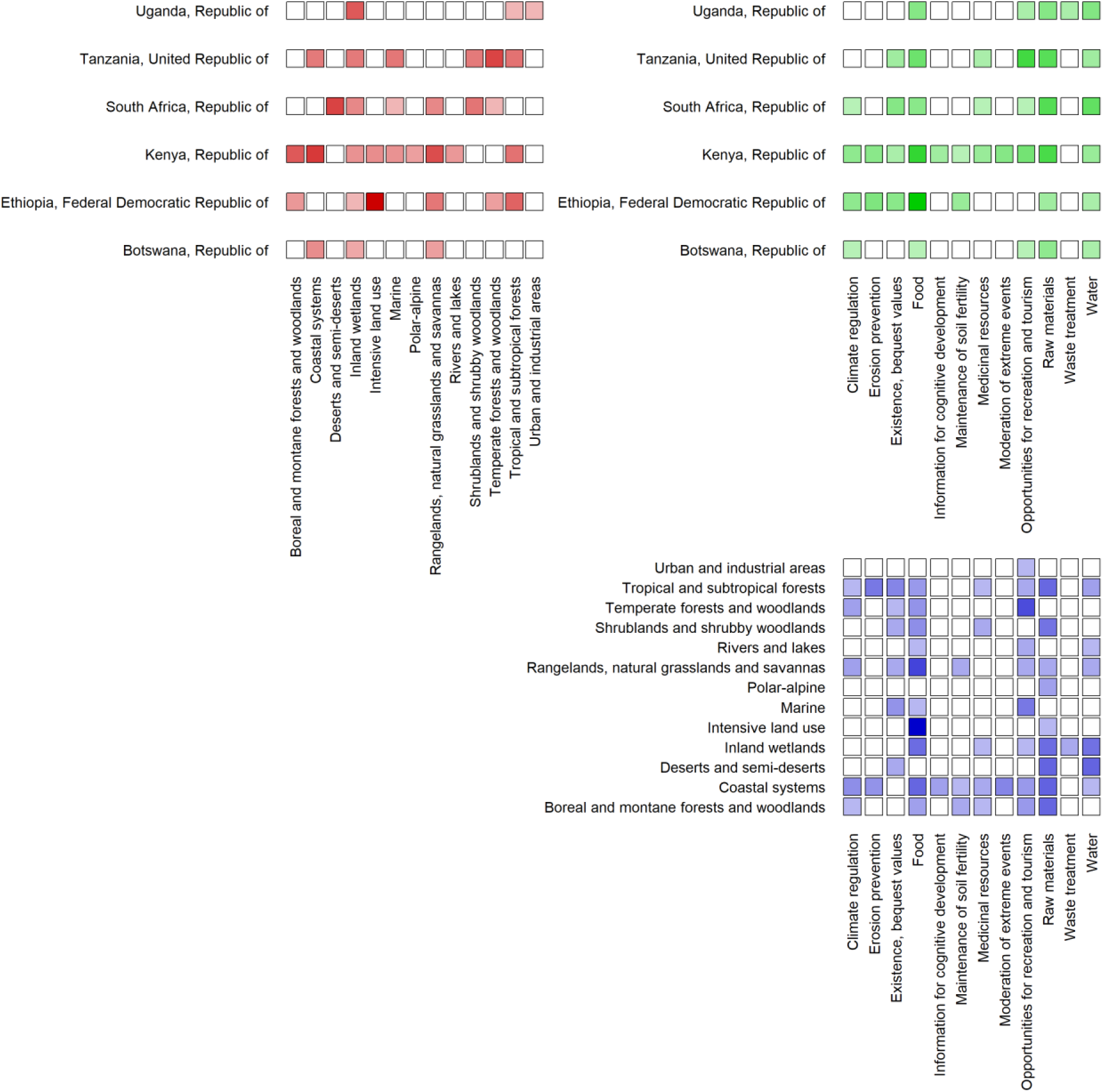
Pairwise combinations of country, type of ecosystem service, and biome for records from the dataset used in our analyses. Here darker shades correspond with more records of estimated ecosystem service value.

**Table S1:**
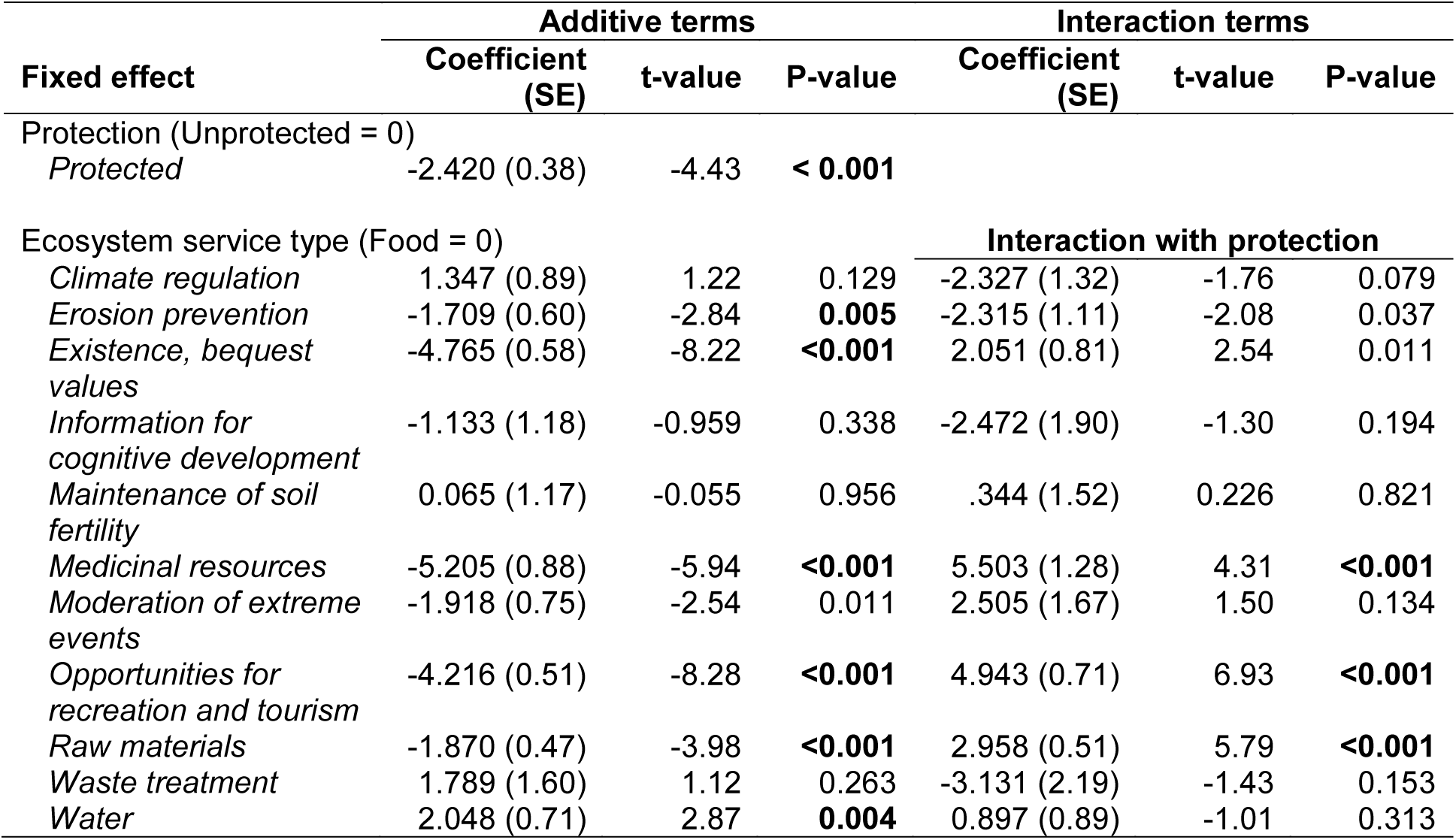
Model coefficients of a log-linked Gamma generalised linear mixed effects model of the standardised value of ecosystem services relative to protected status, the type of ecosystem service and their interaction. Here country and biome are included as random effects in the model. Given the log-link in the model, exponents of the estimated coefficients reflect the statistical effect on the value of ecosystem services. Statistically significant coefficients at the conservative threshold of *α* = 0.01 are shown as P-values in bold font.

